# Structural constraints acting on the SARS-CoV-2 spike protein reveal limited space for viral adaptation

**DOI:** 10.1101/2025.10.27.684767

**Authors:** James C Herzig, Michael L Magwira, Simon C Lovell

**Affiliations:** Division of Evolution, Infection and Genomics, School of Biological Sciences, University of Manchester, M13 9PT, UK

## Abstract

The SARS-CoV-2 pandemic resulted in an unprecedented scientific response. The enormous scale of global genome sequencing, protein structural determination and targeted studies of protein and variant dynamics has resulted in a unique dataset which provides a valuable resource for understanding of viral evolutionary dynamics. Previous analysis of SARS-CoV-2 evolution has revealed apparently saltatory dynamics, with viral variants arising following large evolutionary jumps without genetic intermediates represented in the sequence database. We utilise rich SARS-CoV-2 datasets to interrogate the role of protein structural constraint in SARS-CoV-2 evolution and whether these evolutionary jumps may result from the viral spike protein accessing new regions of viable sequence space. We apply multiple computational predictors of structural constraint across different structural backgrounds and assess how constraint has changed during SARS-CoV-2 variant evolution. These predictions are validated using substitution data from the SARS-CoV-2 global sequence database. We find that all predictive methods suggest that the structural constraint experienced by specific sites has undergone very limited change, despite significant phenotypic evolution of the SARS-CoV-2 S protein. Signature mutations for variants of concern are not found to be under structural constraint by any computational predictor regardless of which viral variant structure is used to calculate predictions. We also develop a machine learning model to assess substitution viability, combining predictors of evolutionary constraint with information about local structural context. This confirms our conclusions, with model performance largely unaffected by the use of different viral variant structures. We also find no reduction in the shared proportion of accessible substitutions over evolutionary time, as would be expected if the S protein had entered and explored novel sequence space during variant evolution. These results suggest that despite its rapid rate of mutation, the SARS-CoV-2 S protein is subject to strict structural constraints and shows that viral genomes exhibit limited plasticity following infection of a new host.

## Introduction

The SARS-CoV-2 pandemic was the worst pandemic of an infectious disease in recent decades, causing global mortality, economic damage and social disruption. However, the response to the pandemic using contemporary technologies such as affordable mass sequencing has resulted in an unprecedented scientific dataset (Hill et al. 2023; Ladner and Sahl 2023). The quantity and temporal granularity of sequences available in SARS-CoV-2 global genome database, and the large number of viral protein structures solved from the earliest period of the pandemic (Arya et al. 2021) provides a unique opportunity to study viral evolutionary dynamics and any evolutionary constraints that may arise form protein structure.

SARS-CoV-2 has undergone several distinct phases of phenotypic evolution. An initial period of largely neutral diversification ended in late 2020 when multi-mutant variants began to arise (MacLean et al. 2020; Harvey et al. 2021; Martin et al. 2021). Those variants with suspected phenotypic characteristics, such as increased transmissibility or immune escape properties, were classified as variants of concern (VOCs) by the World Health Organisation (WHO). The first VOCs were largely characterised by mutations in the SARS-CoV-2 spike (S) attachment glycoprotein associated with an increase in transmissibility (Davies et al. 2021; Campbell et al. 2021; Carabelli et al. 2023). In late 2021, a new VOC given the WHO designation Omicron arose, which was primarily characterised by greater antigenic distance from preceding variants and escape from natural and vaccine mediated immunity (Willett et al. 2022; Cao et al. 2022; Cao et al. 2023; Zhou et al. 2023).

SARS-CoV-2 VOCs with phenotypically distinct characteristics have arisen through apparently saltatory evolutionary dynamics, with novel variants rooted in ancestral sequences that are no longer predominant at the time of variant emergence and limited sequence coverage of intermediate genomes (Viana et al. 2022; Hill et al. 2022). All four major VOCs that emerged from late 2020 to early 2022 exhibit this property (Roemer et al. 2023). This characteristic appears to be due to intrahost viral evolution during chronic infections, in which relaxed selective environments and superinfections result in novel combinations of mutations, with phenotypic properties arising that could not readily arise during typical transmission (Kemp et al. 2021; Wilkinson et al. 2022; Harari et al. 2022). This raises the question of whether the acquisition of novel combinations of mutations distinctive of most VOCs to date occurs due to viral proteins accessing new areas of thermodynamically permitted sequence space, thereby permitting the acquisition of previously non-viable mutations with beneficial fitness effects.

Here, we assess the constraint experienced by the SARS-CoV-2 S protein in order to test whether the sequence space accessible to the S protein changed during variant evolution. Our analysis is enabled by the high sequence coverage and high-quality structures of viral variant S proteins. Many studies have characterised individual mutations (e.g. Jangra et al. 2021; Liu et al. 2022; Lubinski et al. 2022), particularly in the receptor binding domain (RBD), but there has not been any systematic exploration of SARS-CoV-2 S protein structural constraints across different viral variants. We carried out this structural stability-focused analysis to assess the general importance of these constraints to SARS-CoV-2 evolutionary dynamics. We employ multiple computational predictors of constraint, conducting computational deep scans to assess the likely impact of every possible substitution across the SARS-CoV-2 S protein with each method. We also developed a supervised machine-learning model using these predictors and additional features to explore whether combining different predictive methods improved predictions.

Despite the unprecedentedly rich and granular dataset, we find no evidence that structural constraints have changed substantially or played a major role in the acquisition of new mutations during SARS-CoV-2 S protein variant evolution. There is limited evidence that the viral spike protein has acquired any substitutions that were not viable in the Wuhan wildtype. This work therefore supports the view that despite high mutation rates and short-term evolutionary responsiveness, RNA viruses experience strong constraints and are limited to sampling from a limited set of substitutions. This has implications for the evolvability of RNA viruses following spillover into new hosts and the forces likely to drive the macroevolution of coronaviruses.

## Results

SARS-CoV-2 evolution is constrained by the necessity of maintaining thermodynamically stable and biologically functional proteins. To elucidate the importance of structural constraint to SARS-CoV-2 variant evolution, we used computational predictors of constraint to define an envelope of non-viable sequence space. We calculated several site-independent predictors of structural and functional constraint. Briefly, these predictors were:

1. Position specific scoring matrices (PSSMs), providing a measure of phylogenetically derived constraint based on substitution patterns in related sequences, calculated using DELTA-BLAST (Boratyn et al. 2012).
2. Environment specific substitution tables (ESSTs), providing a measure of constraint based on the local structural environment (Overington et al. 1992).
3. Empirical forcefield simulations with FoldX and Rosetta, providing estimates of the impact of substitutions on protein thermodynamic stability (Guerois et al. 2002; Alford et al. 2017).

These predictors were selected to capture constraints arising from the necessity of maintaining a stable, folded protein. In order to explore the spatial and temporal changes to this envelope of inaccessible sequence space we examined predictions made by the constraint predictors across different periods of SARS-CoV-2 evolution. We validated predictions using binary classification of substitutions as either deleterious (occurring below the mean frequency) or non-deleterious (occurring above the mean frequency). This classification was based on the proportional frequency of substitutions at any given site in the GISAID global database of SARS-CoV-2 sequences (Khare et al. 2021).

### Assessing structural constraint across multiple predictors

We made predictions using the four predictive methods described above for every possible substitution across four S protein structures. These were a wild type (WT) structure (PDB: 6VXX), and three VOC S protein structures (Alpha, PDB: 7LWS; Delta, PDB: 7V7N; Omicron, PDB: 7WP9), allowing us to identify differences in the structural permissibility of mutations on different epistatic backgrounds. To ensure the sequence data used to validate predictions was appropriate to these epistatic backgrounds, we used sequence data from four distinct periods for each variant structure, broadly aligned with the period of initial emergence and regional or global predominance for each variant. Results of the computational deep scan are shown in Figure 1, with points showing the predicted constraint for every possible substitution and colours showing the substitution classification as low or high frequency according to observed sequence data. VOC substitutions are highlighted. Note that for FoldX ΔΔG positive scores imply a substitution is more destabilising, while all other scores are positive correlated, with negative scores implying greater destabilisation.

**Figure 1:**
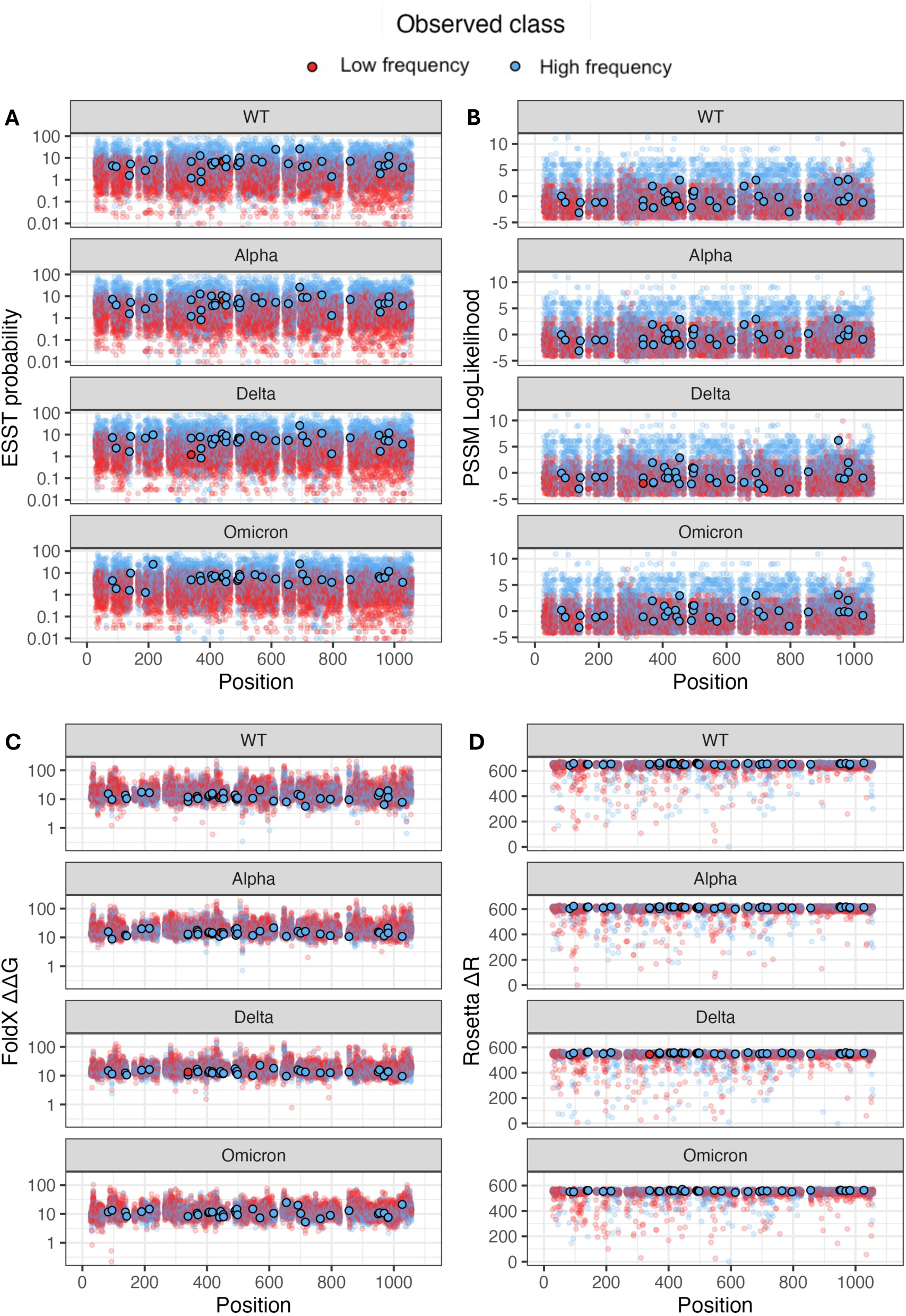
Distribution of constraint predictor scores on the SARS-CoV-2 S protein. Distribution of scores are shown for PSSM log likelihoods **(A)**, ESST probability **(B)**, FoldX ΔΔG **(C)** and Rosetta ΔR **(D)**. Putatively functional substitutions observed in VOCs and their reversions are highlighted. Points are coloured by their classification in observed SARS-CoV-2 sequences deposited in the 6-month period covering the emergence and initial predominance of each variant. Sequences used to validate WT predictions were drawn from the 6 months immediately preceding the emergence of the first VOCs. FoldX and Rosetta scores are normalised preserving rank order to ensure all scores are positive. FoldX and ESST scores are shown on a log scale for clarity. **Alt text:** Scatter plots showing the results of computational deep scans with four predictors of structural constraint, with subfigures labelled A-D.

Both ESST and PSSM predictions (Figure 1A and 1B) show a clear correlation between predictor score and observed substitution class, with the great majority of substitutions that are predicted as highly probable by these methods classified as high frequency based on the sequence data. Predictors based on empirical force fields did not show a similar clear correlation between predicted scores and representation in the sequence database. Plotted VOC substitutions generally have moderate predicted constraint scores by all methods across all epistatic backgrounds, with no VOC substitutions predicted as being under strong structural constraint even in the context of the WT structure, suggesting these substitutions were structurally viable immediately following SARS-CoV-2’s jump into the human population. To test whether there was any significant difference between VOC substitutions and the general case, we performed Mann-Whitney U tests with the hypothesis that the mean value of each constraint predictor was greater or lesser in VOC substitutions relative to the full set of substitutions. VOC substitutions were found to be under significantly less constraint than the mean value across the entire deep scan, as predicted by PSSM, ESST and FoldX (p = 0.0010, 0.000012, 0.034 respectively), but not Rosetta predictions. However, significance was lost when VOC substitutions were instead compared to only substitutions classified as high frequency in the observed data. This suggests that the significant differences reported above are simply reflective of comparing any set of substitutions known to occur in nature with the full deep scan containing every theoretically possible substitution.

All but two VOC substitutions are also classified as high frequency (blue points) according to the thresholding of the sequencing data. This is the case even during the earliest time period sampled (Figure 1A); that is, functional substitutions associated with VOCs were occurring at moderate background levels prior to the emergence of any VOC. Additionally, VOC substitutions associated with the Alpha variant continue to be classified as high frequency and computational predictors do not assess them as destabilising against the background of Omicron variant (Figure 1D) and reversions of VOC-associated substitutions likewise remain structurally viable in the context of the VOC S protein structures. The only exceptions were K444T, which occurred below the classification threshold in both the wildtype and Alpha dataset, and G339H, which occurred below the classification threshold in the Delta dataset. The former is a key neutralising antibody escape mutation present in many Omicron lineages while the latter is also a putative escape mutation which also enhances viral infectivity known from some Omicron lineages. However, while they occurred below the classification threshold in the sequence database during some periods, these substitutions were not assessed as destabilising by any computational predictor. Their classification as low frequency may therefore reflect the limits of our data labelling strategy rather than be a true identification of these substitutions as deleterious in these specific evolutionary periods.

To ensure we were not underestimating the structural constraint experienced by the S protein due to its ability to undergo conformational change, we carried out an additional computational deep scan using FoldX and ESST predictions on open and closed conformation S protein structure. Figure S1 shows that both methods predict the open conformation structure to be subject to reduced constraint on average, with particularly large differences in the S1 subunit. This suggests that the comparison between viral variant closed conformation structures shown in Figure 1 will capture the majority of structural constraint, despite not assessing the constraint experienced by the S protein in the alternative conformation.

These results suggest that fixation of putatively functional VOC substitutions has not been driven by changes to their structural viability, as almost all of these substitutions are predicted to be structurally viable at the beginning of the pandemic and have remained so across different viral variants, and with few exceptions have also occurred above the mean classification threshold.

However, the limited time periods selected to investigate structural constraint may elide important changes in structural viability over the course of SARS-CoV-2 evolution. In order to investigate the consistency of these results over time we trained a supervised machine learning (ML) model to identify low frequency substitutions. Machine learning should maximise the informativeness of our predictors and allow for detection of context-specific or non-linear relationships between predictors.

### Validating a machine learning model of structural constraint

We next developed a machine learning model to enable detection of higher-order structural constraints that are not fully described by any individual method but emerge through interactions of multiple predictive features. To exploit the potential of a machine learning approach we expanded the feature set, with the full set of predictors used to inform the final ensemble random forest model comprising local structural contextual information, raw energy terms from FoldX and Rosetta, and predicted T cell epitopes in addition to the predictors already assessed in Figure 1. Labels for supervised learning training data were established through the observed proportional frequency of substitutions at every locus in the GISAID sequence database as described above.

We initially trained a model using the wildtype structure and data labels established for the earliest time period used in Figure 1A, prior to the emergence of any variants of concern. Random forest was selected as the optimal algorithm, outperforming GLM-based approaches. Training on 60% of randomly sampled substitutions from the wildtype protein structure and testing on the remaining 40% showed the model had excellent discriminatory power for substitutions occurring in the same structural and temporal context (Figure 2A). We then re-trained the model on the wildtype structure using all substitutions and tested its performance on the three variant structures using the same data labels as applied in Figure 1. We found that the model was able to correctly assign the substitution viability for the majority of possible substitutions with areas under the ROC curve (ROC AUC) of 0.81, 0.8 and 0.78 on the Alpha, Delta and Omicron S proteins respectively (Figure 2B). In order to gauge the success of our data labelling strategy, we also trained the model using experimental deep scan data on the SARS-CoV-2 wildtype RBD as data labels (Starr et al. 2022). Deep scan protein expression data was used to generate binary classes in the same manner as substitution frequencies. Models trained to predict these data labels achieved superior performance to those using substitution frequency data labels, with ROC AUC values of 0.93, 0.92, 0.83 for Alpha, Delta and Omicron S protein respectively, compared to values of 0.86, 0.84 and 0.79 using substitution frequency data labels (Figure 2C). While the superior performance indicates there is a stronger relationship between computational predictors and the experimentally-derived labels, the limitation of deep scan data to the RBD and wild type structure limits their value in training models applicable to the entire S protein.

**Figure 2.**
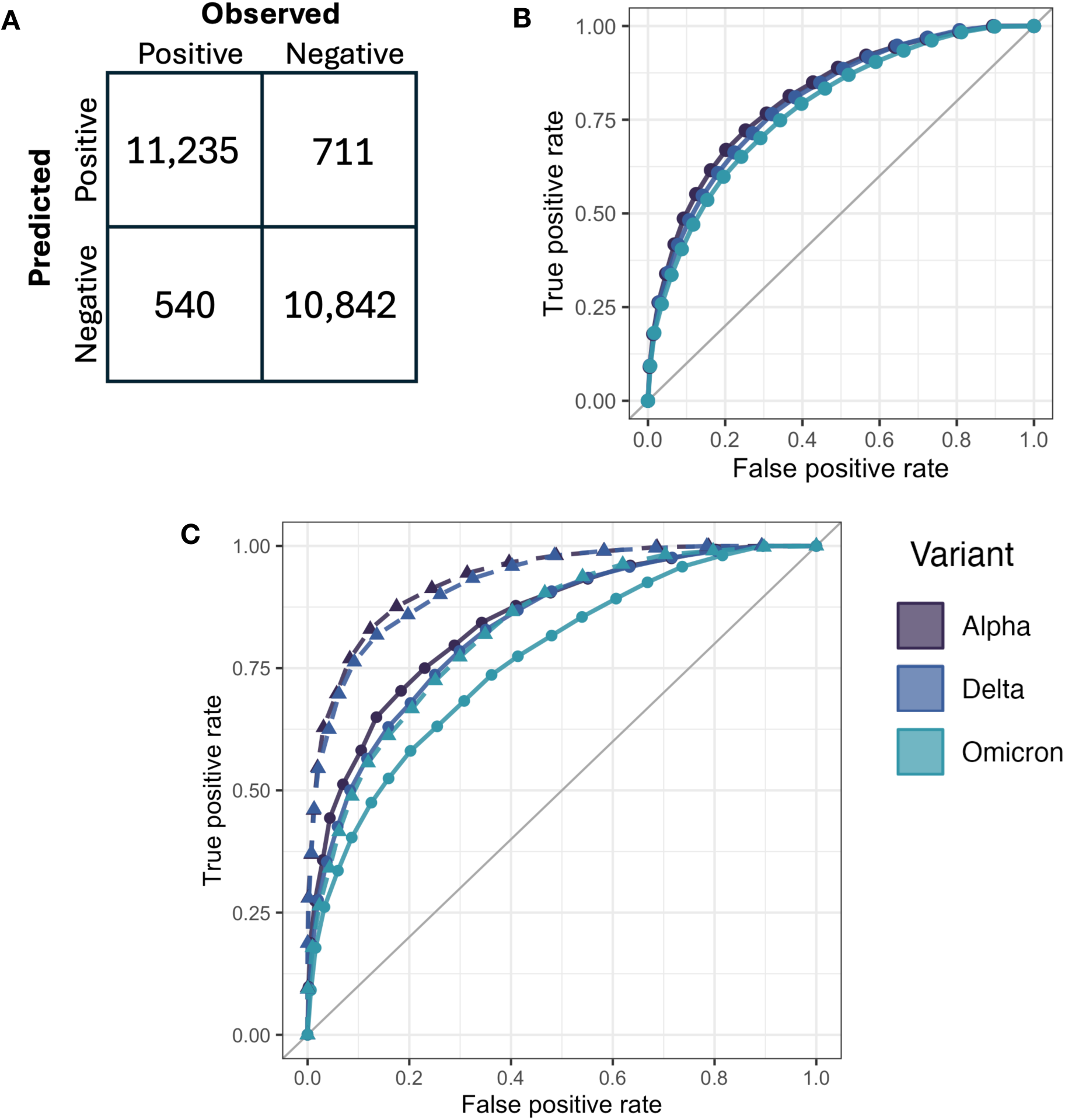
**A** shows the confusion matrix for the validation set of substitutions from the wild-type dataset. A random forest model was trained on 60% of substitutions from this dataset and tested on the remaining 40%. **B** shows the ROC curves resulting from application of this model to the Alpha, Delta and Omicron datasets. **C** shows performance of the model in the RBD region only when trained with data labels based on frequency in the GISAID database (solid lines) or data labels based on experimental deep scan data (dashed lines). **Alt text:** Graphs and figures showing the performance validation of the supervised machine learning model when predicting S protein substitution viability for three VOC structures, with subfigures labelled A through C.

Overall, the model performed well and was able to correctly predict the frequency of substitutions in most cases. Performance of model predictions did decline slightly over evolutionary time, with predictions made on the Omicron structure and data labels showing somewhat weaker performance, although performance between variants was very similar across the whole S protein (Figure 2B). We therefore aimed to conduct a more thorough and granular analysis to identify whether this reflects a change in structural constraints over S protein evolution. By training models on all variant structures and using data labels calculated from across the entire initial period of SARS-CoV-2 variant diversification we aim to explore to what degree structural constraints changed during this period and how they may have influenced variant evolution.

### Predicting structural constraint across SARS-CoV-2 variant evolution

To ensure the substitution viability labels reflected more granular changes during variant evolution, cut-off dates were set at the 1^st^ of every calendar month from December 2020 – December 2022 and models were retrained at every cut-off date. Training sets were sampled from all sequences deposited in the 3-month period prior to each cut-off date and test sets used all sequences deposited in the 3-month period subsequent to each cut-off date. Although no sequences appear in both training and test datasets, the majority of substitutions do not change class between the training and test periods, resulting in many identical feature/outcome pairs. To alleviate over-fitting to this majority case, we applied an ensemble model, sampling the training data to generate 10 subsamples, training models on each subsample and then establishing a final prediction based on majority vote of the individual models.

Aggregate performance of models trained using predictions calculated with wild type and variant structures is shown in Figure 3A. The area under the ROC curve is shown for models trained at every cut-off date using each of the four protein structures. The ML model correctly predicted substitutions that maintained their class between train and test sets in all cases, resulting in the strong aggregate model performance shown in Figure 3A. However, the minority case of substitutions which change class between consecutive training and test sets, hereafter referred to as ‘class-switching substitutions’, were mostly incorrectly classified by our structural constraint-based model (Figure S2). To identify whether this was due to these sites being under selection or undergoing changes to their thermodynamic viability between training and test sets we assessed the relative change in frequency for these class-switching substitutions between every consecutive training set. We found no statistically significant difference between the mean or median change in substitution frequency for these positions (paired T test, median p = 0.22, mean p = 0.44). These substitutions are not changing frequency at a greater rate than the general population and are therefore likely caused by stochastic movement around the classification threshold rather than genuine changes in their thermodynamic viability.

**Figure 3.**
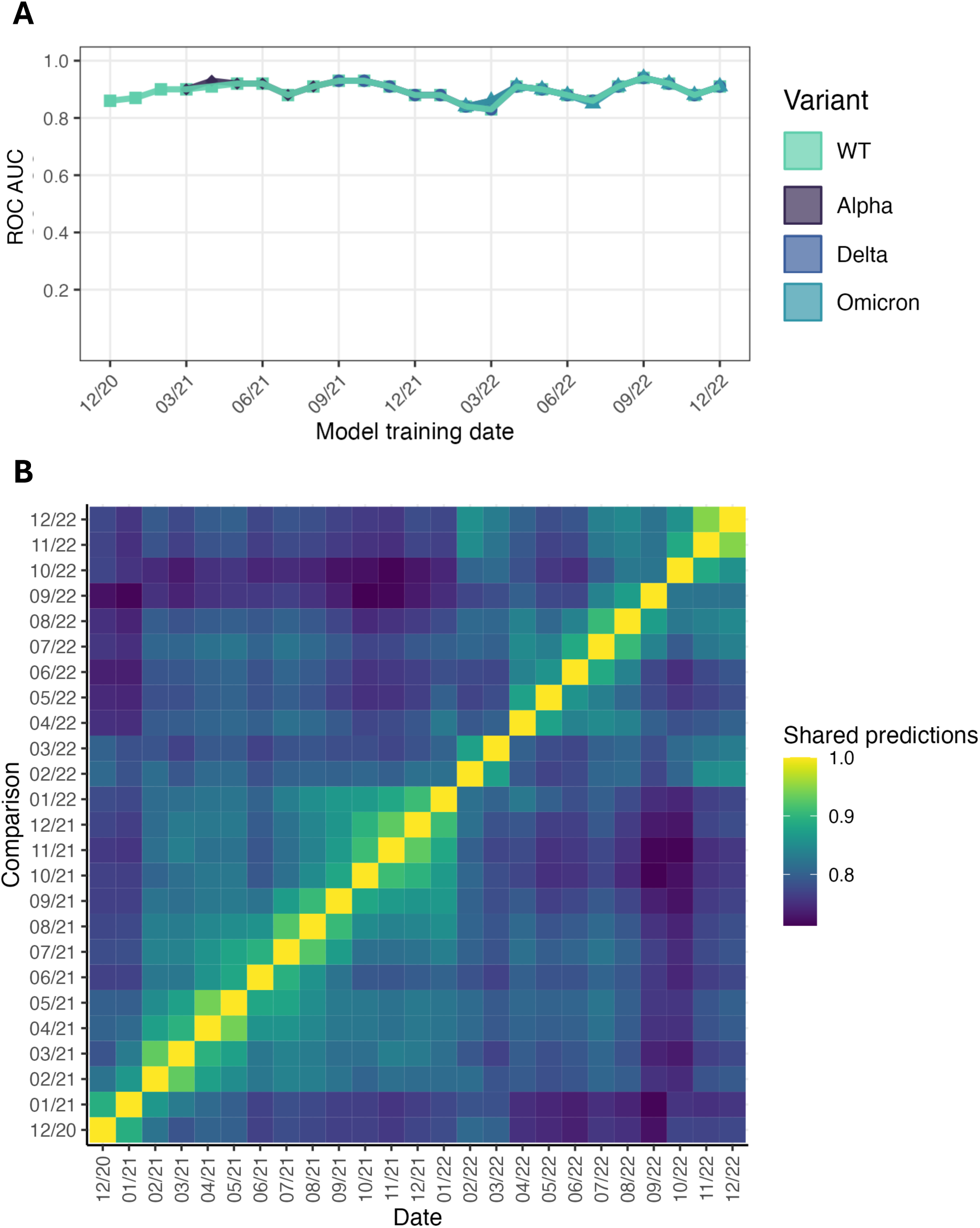
**A** shows the outcomes of ML predictions of substitution class. Notably, the use of S protein variant structures resulted in almost no difference in model performance. **B** shows the proportion of substitutions with the same predicted classification between every time period studied, calculated using the WT structure. **Alt text:** Graphs and figures showing the performance of the supervised machine learning model when predicting S protein substitution viability in different temporal and structural contexts, with subfigures labelled A and B.

Both aggregate performance and performance on class-switching substitutions was nearly identical regardless of the S protein structure used, suggesting that structural constraint did not substantially change between different S protein VOCs. However, similar performance does not necessarily imply that the specific set of viable substitutions remained identical over time. We therefore assessed the proportion of substitutions that share the same model classification between every studied time-point. Results of this analysis are shown in Figure 3B and reveal a temporally local increase in the proportion of substitutions with shared classifications, but no long-term directional change. This pattern was maintained regardless of the variant input structure used to calculate constraint predictors.

Finally, to identify effects of the length of the sampling period, we repeated modelling using test sets derived from 1-,6- and 12-months of sequence data subsequent to the cut-off date. Performance over time was relatively consistent (see Figure S3). Shorter test periods, which contain sequences with a lower average evolutionary distance from the training set, resulted in slightly superior but more erratic performance. This is reflected in the classification of VOC substitutions, with sampling of sequences from shorter time periods resulting in greater stochasticity, as shown in Figure S4. Longer sampling periods result in almost all VOC substitutions being classified as high frequency, as shown in Figure 1, but 1- and 3-month test samples resulted in numerous VOC substitutions being classified as low frequency for short periods. This short-term stochasticity in substitution frequency increases confidence that variability in our model predictions is reflective of variability in the data labels due to the mixed, complex fitness effects of many substitutions or phylogenetic factors rather than any genuine change to structural viability.

### Identifying sites with shifting constraints

We have established that the majority of positions have maintained similar degrees of structural constraint over the course of SARS-CoV-2 variant evolution, including positions implicated in phenotypic changes in S protein immunogenicity and antigenicity. We now aimed to identify any outlying sites at which structural constraints had changed by inspecting those substitutions for which model predictions changed over the period of variant evolution. Conducting a manual assessment of all of these substitutions is not feasible; we therefore conducted further filtering to identify any of these substitutions which underwent particularly large or consistent changes in frequency, which may indicate changes to their thermodynamic viability following the rapid expansion of a new viral lineage. We filtered identified substitutions with three criteria: those that had undergone consistent changes in frequency over six consecutive months, those which underwent particularly large changes between any two consecutive training periods (97.5^th^ percentile) and those which reached a frequency > 0.8% in the overall population of sequenced virus at any time during the studied period. We identified 118 substitutions at 103 unique sites with these criteria to analyse in more depth. We assessed the identified substitutions by inspecting the outputs of individual constraint predictors, their observed frequency over time in the sequence database and the local environment of the substitution in wild-type and variant protein structures. Through this process we identified only a single amino acid substitution, C336H, for which all assessed data suggested it may have been enabled by changes to local structural constraints during SARS-CoV-2 variant evolution. C336 forms a disulphide bond with the cysteine at position 361 (Figure 4A) and replacement of C336 by histidine is predicted as highly thermodynamically unfavourable by both Rosetta and FoldX simulations in the wild-type structure.

**Figure 4.**
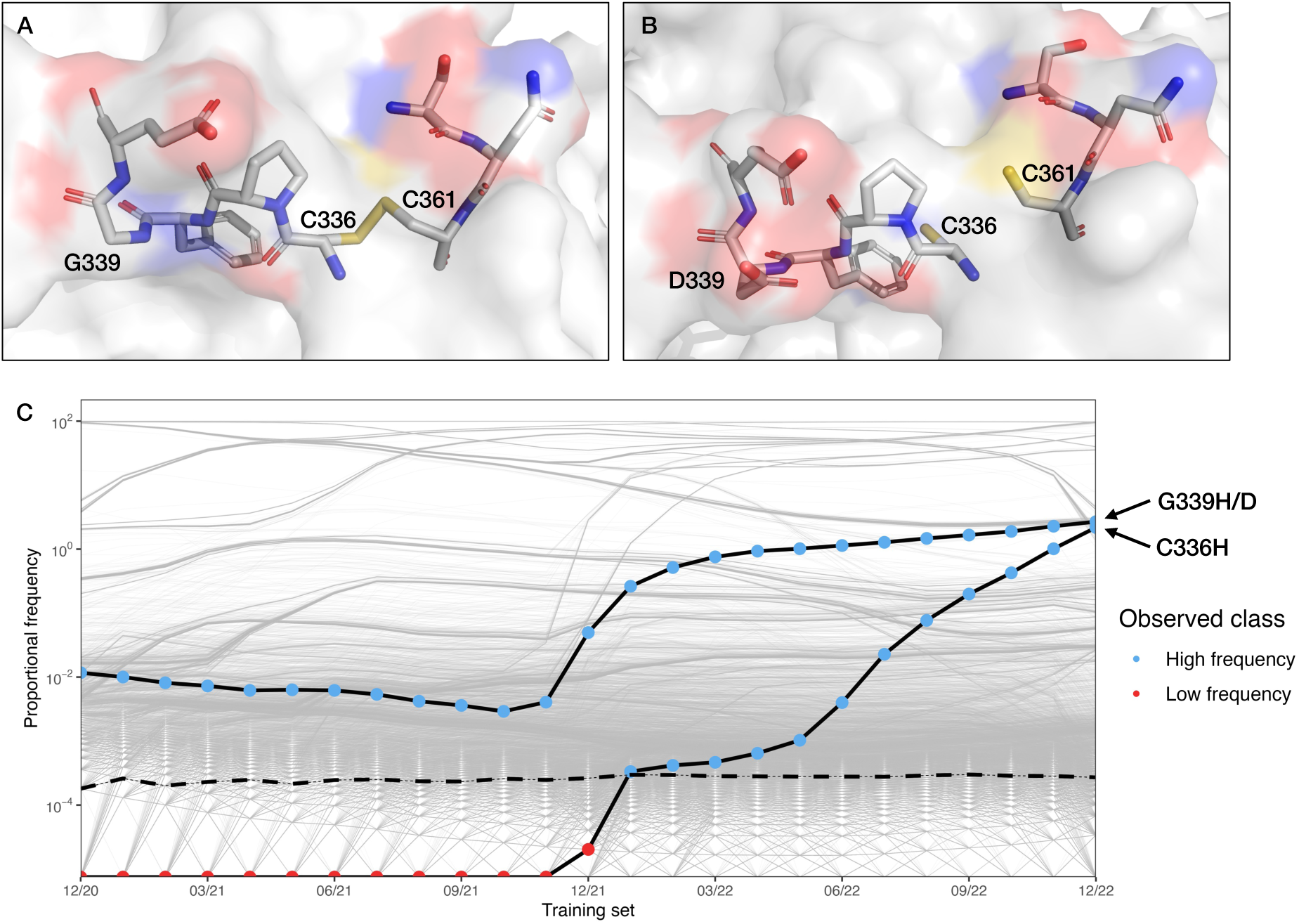
**A** shows the local structure of C336 in the SARS-CoV-2 wild-type S protein (PDB: 6VXX). **B** shows the local structure of C336 in the SARS-CoV-2 Omicron S protein (PDB: 7WP9). Colours show oxygen (blue), nitrogen (red) and sulphur (yellow) atoms. The loss of the disulphide bond and changes to surface charge caused by the G339D substitution can be seen in the Omicron S protein structure. **C** shows the observed frequency of C336H and G339H/D in the sequence database. The points are coloured to show the observed class of the substitution in each training dataset. The classification threshold is shown by a dashed line. Frequency of all other substitutions are shown by pale grey lines. **Alt text:** Figure showing the local molecular structure of the SARS-CoV-2 S protein proximal to residue C336 and a graph showing the frequency of C336 and G339H/D in the sequence database over time, with subfigures labelled A, B and C.

However, in the Omicron structure Rosetta, FoldX and ESST predict a large reduction in the destabilising effect of these substitutions relative to the wildtype, although it is still assessed as destabilising by FoldX (ΔΔG: 73.9 → 21.9; ΔR: -4.7 → 17.3; ESST probability: 0.24 → 1.26). Many Omicron lineages feature the G339H/D substitution which has been proposed to contribute to the Omicron S protein’s improved infectivity (Pastorio et al. 2022), with evidence suggesting the increased surface charge plays a role in altering immunogenicity (Zhang et al. 2022, Figure 4B). It is plausible that replacing the glycine at position 339 with an amino acid with a charged side chain may partially alleviate the cost to protein stability of a histidine substitution at position 336, enabling the C336H replacement to occur. This conclusion is supported by the observed frequency of this substitution during variant evolution. C336H was not detected in any sequences until December 2021, at which point it rapidly increased in frequency (Figure 4C). This increase tracks the increase of Omicron lineages generally and the G339H/D substitution specifically. This data therefore supports, although does not confirm, that the H336 was stabilised by structural changes in the Omicron S protein, enabling the substitution to arise through changes to structural constraint.

## Discussion

We have found no evidence that changes to the thermodynamically viable sequence space of the S protein contributed to the phenotypic evolution of SARS-CoV-2 variants. The predictions of individual computational methods for VOC mutations (Figure 1) and aggregate ML model performance (Figure 3A) were consistent across different epistatic backgrounds. Furthermore, the proportion of substitutions with shared classification showed no directional temporal change (Figure 3B), as would be expected if the S protein was moving through sequence space and sampling previously non-viable substitutions. Despite the strong selective pressure acting on the SARS-CoV-2 S protein to increase antigenic distance and avoid host immune responses (Willett et al. 2022; Cao et al. 2022) there has been no decrease in the shared set of viable substitutions over time; this strongly suggests that the SARS-CoV-2 S protein is subject to hard structural constraints that cannot easily be overcome over the evolutionary short term.

Our approach of assigning substitution viability by thresholding of the proportional frequency of substitutions as represented in the global genome sequence database will result in some inherent variance and incorrect model predictions. Significant bias is known to exist in the global sequencing database (Wohl et al. 2023) and the SARS-CoV-2 S protein has positions under antagonistic pleiotropy (e.g. those with differing inter- and intra-host fitness effects; Lee et al. 2023), although it has been shown that only a small fraction of common intra-host single nucleotide variants rise to a high frequency and therefore their effect is likely to be limited (Hou et al. 2023). The impact of such biases is challenging to quantify, but it is clear that a substitution’s frequency in the global database is not necessarily a true reflection of its viability in all cases. The improved performance of RBD predictions made using data labels based on experimental deep scan data (Figure 2C) suggests that some information in the predictive computational features is lost due to noise present in the data labels based on substitution frequencies. Nonetheless, the clear correlation between predicted ESST and PSSM scores and classifications based on the sequence data in Figure 1 suggest that despite these problems, a binary classification approach is able to distinguish at least strongly deleterious mutations. The majority of proteins can tolerate only minor decreases in stability before the fraction of partially or mis-folded protein causes a significant fitness cost and loss of the mutant variant (Yue et al. 2005; Bershtein et al. 2006). Substitutions with substantial negative effect on thermodynamic stability are therefore likely to be highly deleterious and will be correctly classified as low frequency using this thresholding approach. Meanwhile, mutations with more complex fitness effects may show greater susceptibility to short-term fluctuations in viral lineage abundance and sequencing bias, as shown by the changing classifications of some VOC substitutions in Figure S4.

Inspecting sites where model predictions changed over time identified one site where changes to thermodynamic viability may have enabled a novel substitution to arise. Our analysis of the S protein structure, C336H’s frequency over time and the outputs of our computational predictors suggest that changes to the Omicron S protein structure may be prerequisite to substitution of the cysteine at position 336. The directionality of this relationship is revealed by the fact that C336H is not present in many Omicron sequences carrying G339H/D; C336H is evidently not a compensatory stabilising mutation necessary for the fixation of G339H/D. Therefore, this site does not alter our conclusion that changes to structural constraints have played a very limited role in the phenotypic evolution of the S protein. Indeed, it appears that the inverse may have occurred: selection of a substitution with beneficial effects on S protein infectivity and immunogenicity enabled the only novel substitution we can ascribe to altered structural constraint with any confidence.

Our conclusion that the set of structurally viable substitutions has not substantially changed during the period of rapid SARS-CoV-2 variant evolution suggests that phenotypic evolution is instead driven by novel combinations of mutations with functional epistatic relationships. This is supported by evidence from experimental deep scans that has found strong functional epistasis between sites in the RBD involved in binding to the ACE2 receptor resulting in greatly increased receptor affinity in combination, but not individually (Starr et al. 2022; Moulana et al. 2022). Rosetta modelling carried out by Rochman et al. (2022) also supports our results, finding that the structural constraint acting on the RBD has been maintained across all VOCs and that the escape mutations present in Omicron strains likely represent the majority of structurally-viable substitutions in the RBD. We therefore propose that emergence of SARS-CoV-2 variants occurred through novel combinations of substitutions that were sampled from a limited region of structurally viable sequence space. Complex fitness effects and functional epistatic relationships result in fitness valleys that prevent stepwise acquisition of these substitutions in the course of typical transmission and evolution, regardless of their non-deleterious effects on protein thermodynamic stability. Relaxed constraints during chronic infections provide opportunities for the virus to cross these fitness valleys and acquire assortments of substitutions which result in new phenotypic properties (Lawrence et al. 2021; Smith and Ashby 2023; Nielsen et al. 2023). However, these relaxed constraints are insufficient to enable escape of the protein from a region of sequence space strictly limited by the necessity of maintaining thermodynamic stability. The SARS-CoV-2 S protein sits on a rugged fitness plateau surrounded by steep sides that cannot be crossed in the evolutionary short term.

This conclusion contributes more broadly to the view that, despite their high rates of mutation and ability to respond rapidly to evolutionary pressures, RNA viruses are equally defined by their genomic fragility which limits longer term evolvability (Belshaw et al. 2008; Sanjuán 2010). Despite SARS-CoV-2’s very large population size, rapid rate of mutation and a selective landscape that underwent major changes following initial zoonotic spill-over, phenotypic change during the period of rapid viral variant emergence was driven by sampling of mutations from a limited set of substitutions which were structurally viable in the viral genotype that initially established infection in the human population. Where substitutions that were non-viable in the wild-type have occurred (C336H), they are downstream of changes to local RBD structure driven by phenotypic evolution of the Omicron variant. This suggests that over longer evolutionary time, the set of viable substitutions may shift; however, this is a slow and limited process even under conditions of host switching and subsequent rapid evolution. The SARS-CoV-2 S protein’s inability to access novel regions of sequence space suggests that RNA viruses are subject to strict constraints that limit their genomic plasticity. Larger evolutionary shifts will instead likely depend on genomic reorganisation or transfers of genetic information, events which have been identified as the major drivers of viral macroevolution (Kitchen et al. 2011; Shi et al. 2016; Koonin et al. 2022).

## Methods

### VOC substitutions

VOC substitutions studied were those addressed by Carabelli et al. (2023) and Harvey et al. (2021). These are substitutions that have been described as impacting SARS-CoV-2 infectivity or transmissibility; they may also contribute to immune escape properties.

### Constraint predictors

#### 2.5.2a PSSMs

PSSMs were generated using DELTA-BLAST as described in the BLAST manual (National Center for Biotechnology Information (US) 2021). All DELTA-BLAST searches were performed locally using the BLAST+ software. Reference FASTA files for structures were downloaded for relevant structures from the PDB and used to generate PSSMs. PSSMs were generated with a single iteration of DELTA-BLAST.

#### 2.5.2b ESSTs

Environment Specific Substitution Table (ESST) (Overington et al. 1992) probability values were calculated as described in Chelliah et al. (2004) after local structural environments were calculated for relevant PDB files using the JOY server (Mizuguchi et al. 1998).

#### 2.5.2c FoldX

Structures were first repaired using the FoldX RepairPDB function with standard settings, which identifies and re-packs the sidechains of residues with high total energy due to e.g. bad torsion angles or stearic clashes. Substitutions were then performed using the repaired structures on all three chains simultaneously using the *BuildModel* function, giving the change in free energy of unfolding, with negative values implying stabilising mutations. FoldX version 5 was used for all simulations

#### 2.5.2d Rosetta

Rosetta simulations were performed using the ‘soft’ repulsive potential functions. Soft-repulsive potential scales van-der-Waals interactions to shorter distances, permitting shorter distance interactions between atoms without large clash scores. For more information, see Kellogg et al. (2011). The structure was first relaxed using the Rosetta Relax function with a fixed backbone, as described in Kellogg et al. (2011). Mutations were then performed using the Fixbb function with all sidechains optimised through repacking of sidechain rotamers followed by optimisation of sidechain torsion angles through minimisation. A final score was obtained by calculating the difference in arbitrary Rosetta Energy Units between the wild-type and mutated structure.

#### T cell epitope prediction

A list of predicted SARS-CoV-2 T cell binding epitopes calculated by the netMHCpan1 software (Hoof et al. 2009) was taken from Saini et al. (2021). The sequence of the relevant S protein structure was searched for matching epitopes and all positions in a predicted binding epitope were flagged to give a binary feature.

### SARS-CoV-2 protein structures

High-quality structures of the SARS-CoV-2 S protein were identified by a search of the RCSB PDB. Structures with high resolution, low clash scores and low Ramachandran outliers were prioritised.

6VXX was identified and used for all analyses, unless otherwise specified. For analysis of open and closed conformation structures, the structures 6XM0 and 6XM5 were used respectively.

Viral variant structures used were 7LWS (Alpha), 7V7N (Delta) and 7WP9 (Omicron).

### SARS-CoV-2 sequence data and classification thresholds

All SARS-CoV-2 S protein sequences were sourced from the Global Initiative on Sharing Avian Influenza Data (GISAID) database (Khare et al. 2021).

Sequences underwent simple filtering for quality. Sequences with an amino acid length of under 1271 were removed from the dataset; this cutoff was selected as it corresponds to the 25^th^ percentile of sequence lengths in the dataset, preserving the majority of sequences while removing short and fragmentary entries. Sequence entries with >1% Ns were also excluded from the analysis.

#### Classification of substitutions

Sequences were used to generate a set of validated outcomes by calculating a consensus matrix of the frequency of each substitution at every position in the sequence using the *Biostrings* package in R. The proportional contribution of every amino acid at every position was then calculated. The mean proportional frequency was calculated for every study period and this was set as the threshold. Any substitution exceeding the mean value was designated in the negative class, ‘high frequency’, any substitution with a proportion less than or equal to the mean value was designated in the positive class, ‘low frequency’. For the experimental deep scan RBD classification used in Figure 2C, protein expression data was taken from Starr et al. (2022). The mean expression value was set as the threshold and substitutions classified using this threshold as described above.

#### Filtering by deposition date

Sequences deposited within the reported date range (inclusive) were included in these datasets.

The individual constraint predictions described in Figure 1 classified substitutions using thresholds calculated from non-overlapping 6-month periods of sequence depositions. Predictions made using the wildtype structure used sequences deposited between 01/07/2020 and 31/12/2020; this largely precedes the emergence of VOCs, with the first Alpha and Beta sequences identified in late October and not reaching significant proportion of total sequences until January 2021. Predictions made using an Alpha structure used sequences deposited between 01/01/2021 and 30/06/2021; this largely covers the entire period in which Alpha represented a substantial proportion of sequences globally. Predictions made using a Delta structure used sequences deposited between 01/07/2021 and 31/12/2021; this covers the period in which Delta was the globally dominant variant, with the first Omicron sequences detected in late November 2021. Predictions made using an Omicron structure used sequences deposited between 01/01/2022 and 30/06/2022, for which Omicron was the globally dominant lineage.

The test sets in Figure S4 are reported as 1-, 3-, 6- or 12-month test sets. These test sets were generated by filtering for all sequences deposited between the reported training cutoff date and exactly 1, 3, 6 or 12 calendar months subsequent to this date. The exact period of time from which sequences were sourced for test sets is therefore not identical, varying by the number of days of different calendar months.

### Machine learning

Random forest algorithms were implemented using the *ranger* package in R (Wright and Ziegler 2017). Models were trained with 25 repeats of 10-fold cross validation. Hyperparameter tuning was performed via a grid search with a tune length of 8. The modal optimal values of hyperparameter tuning on a limited initial training set were calculated and these values were then used for all future ML modelling. The selected values were mtry = 6, splitrule = ‘extratrees’ and minimum node size = 1.

A final feature set of 30 features was selected by recursive feature elimination, with rank aggregation used to select the final features based on a limited initial training set. The final list of features used with a brief description is shown in Table 1.

**Table 1:** Features used for random forest modelling.

| Feature | Description |
| --- | --- |
| WTAA | Wild-type amino acid |
| ReplacementAA | Replacement amino acid |
| **Predictive methods** |  |
| ESST_probability | ESST predicted probability |
| LogLikelihood | PSSM predicted log likelihood |
| total_score | Rosetta-predicted $\Delta R$ |
| pred_epitope_rank | T-cell epitope ranking determined by NetMHCpan 4.1 software |
| **Local structural environment features** |  |
| Ooi_number | Measure of protein packing density |
| **FoldX energy terms** |  |
| Backbone_Hbond | Contribution of backbone hydrogen bonds |
| Sidechain_Hbond | Contribution of sidechain-sidechain and sidechain-backbone hydrogen bonds |
| Van_der_Waals | Contribution of van der Waals forces |
| Electrostatics | Contribution of electrostatic interactions |
| Solvation_Polar | Penalisation for burying polar groups |
| Solvation_Hydrophobic | Contribution of hydrophobic groups |
| Van_der_Waals_clashes | Penalisation for van der Waals' clashes |
| entropy_sidechain | Entropy cost for fixing the sidechain |
| entropy_mainchain | Entropy cost for fixing the mainchain |
| torsional_clash | Intraresidue van der Waals' torsional clashes |
| backbone_clash | Penalisation for backbone-backbone van der Waals' clashes |
| electrostatic_kon | Electrostatic interactions between molecules in the precomplex |
| energy_ionisation | Contribution of ionisation energy |

**Table 1: Features used for random forest modelling**
| Rosetta energy terms |  |
| --- | --- |
| dslf_ss_dih | Dihedral score in current disulphide |
| fa_atr | Lennard-Jones attractive potential between atoms in different residues |
| fa_dun | Internal energy of sidechain rotamers |
| fa_pair | Statistics-based pairing term, favouring salt-bridges |
| fa_rep | Lennard-Jones repulsive potential between atoms in different residues |
| fa_sol | Lazaridis-Karplus solvation energy |
| hbond_bb_sc | Sidechain-backbone hydrogen bond energy |
| hbond_sc | Sidechain-sidechain hydrogen bond energy |
| p_aa_pp | Probability of amino acid at given $\phi/\psi$ angle |
| ref | Reference energy for each amino acid, used to balance internal energy of amino acid terms |

Samples for the ensemble modelling were generated as follows: consider a multiple sequence alignment of SARS-CoV-2 protein sequences *S* of length *L*. Sequences were read from *S* into a subsample *T_i_* with every *i*th sequence dropped from the subsample. This process was repeated *i* times to give *i* subsamples. All modelling presented herein was carried out with *i* equal to 10, resulting in 10 subsamples. Subsamples were therefore each of length *L* - (*L/i*). These subsamples were then processed to generate 10 consensus matrices.

Individual models were trained using the proportional frequency of amino acid identities from each of these 10 consensus matrices. Final class predictions were calculated by simple majority vote of all 10 models, with hung votes resulting in an indeterminate prediction. However, the final predictions made by ensemble models did not feature any hung votes.

Class-switching substitutions were determined as those that were assigned a different class in any of the 10 training subsamples relative to the test sample.

#### Software and packages

Unless otherwise stated, all analysis was performed in *R* version 4.1.1. Analysis and processing of biological sequences and structures was carried out with *SeqKit2* (Shen et al. 2024) and the *R* packages *biostrings* v2.68.1 and *bio3B* v2.4.4.

## Acknowledgements

We gratefully acknowledge all data contributors, i.e., the Authors and their Originating laboratories responsible for obtaining the specimens, and their Submitting laboratories for generating the genetic sequence and metadata and sharing via the GISAID Initiative, on which this research is based. With thanks to Prof. Jeremy Derrick, Dr Amro Safadi and Dr Mitra Kabir for their valuable advice.

This work was funded by a Biotechnology and Biological Sciences Research Council doctoral training centre award to JCH.

## Data availability

A full list of Epi-Set IDs for all sequences are available in EPI_SET_240929he, available from doi:10.55876/gis8.240G2Ghe

Code and data is available at: https://github.com/jcherzig/sars_cov_2_constraint

## Supplementary figures

**Figure S1:**
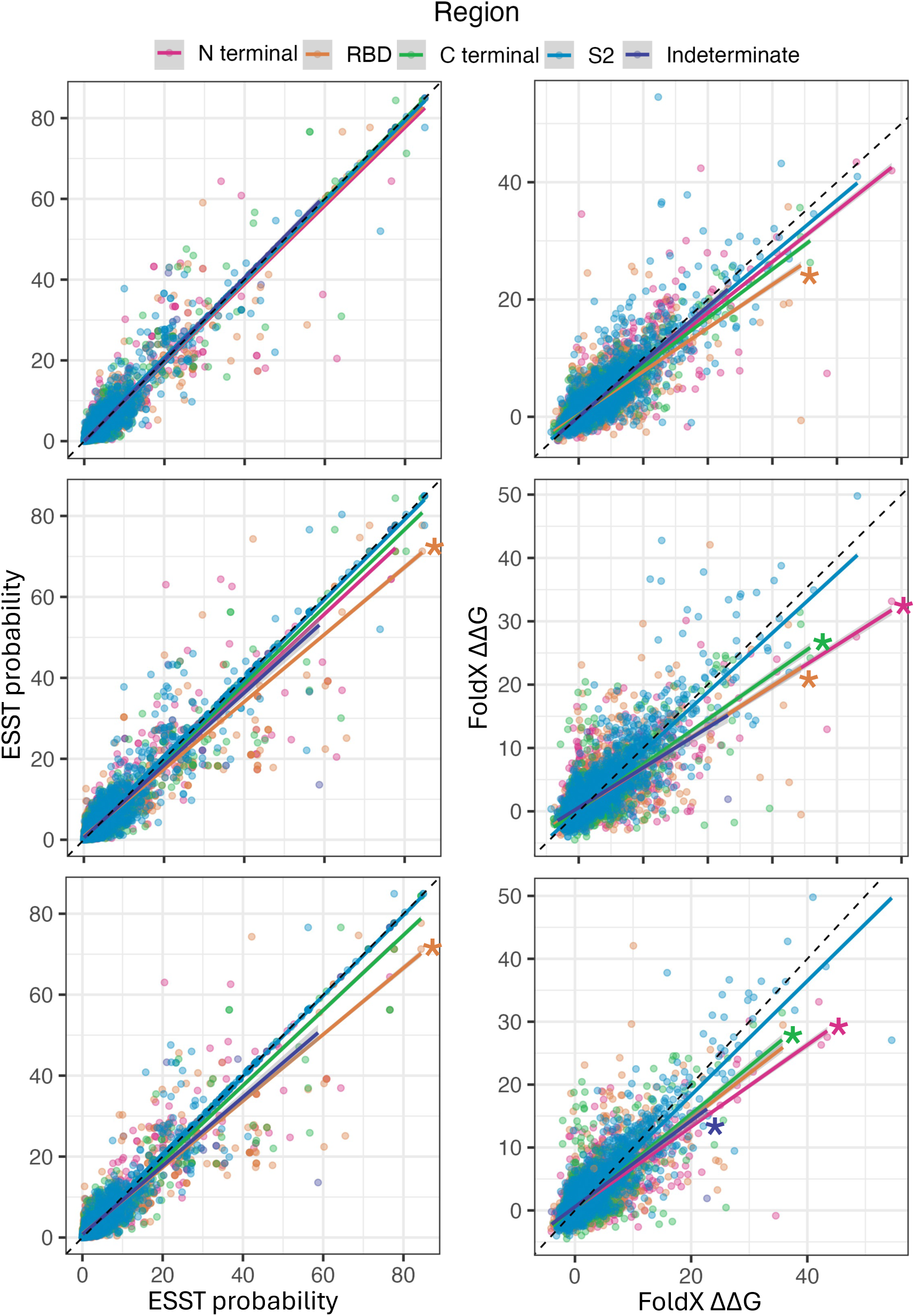
Scatterplots showing correlation of ESST and FoldX scores between ‘up’ and ‘down’ chains in open and closed conformation S protein structures. Scatterplots showing the correlation in ESST (left column) and FoldX (right column) scores between predictions made on open and closed conformation S proteins. The top plots show a closed conformation ‘down’ chain on the x-axis and an open conformation ‘down’ chain on the y-axis; the middle plots show a closed conformation ‘down’ chain on the x-axis and an open conformation ‘up’ chain on the y-axis; the bottom plots show an open conformation ‘down’ chain on the x-axis and an open conformation ‘up’ chain on the y-axis. Substitutions are coloured by protein region. Coloured lines show linear regressions for each protein region. Comparisons with significantly different mean values are marked with coloured asterisks. Dashed black line shows a perfect correlation. **Alt text:** Graphs showing the predicted constraint acting on SARS-CoV-2 S protein in open and closed conformations.

**Figure S2.**
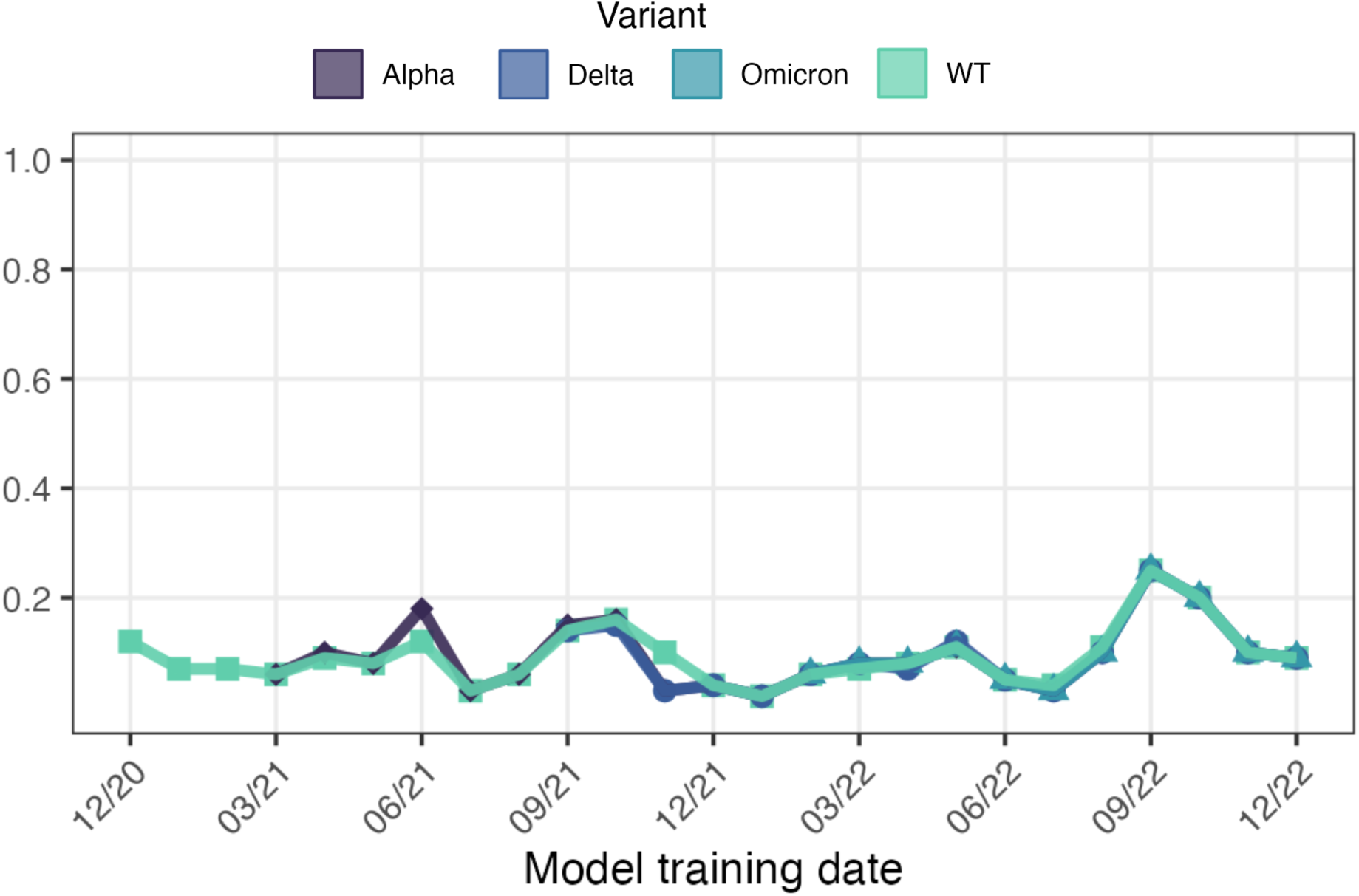
shows the outcomes of ML predictions of substitution class on class-switching substitutions; that is, those that change class between the train and test sets. This accounts for ∼15% of substitutions. The majority of class-switching substitutions are incorrectly classified, resulting in an ROC below 0.5 when these substitutions are analysed independently. **Alt text:** Graph showing performance of the model on class-switching substitutions under different training conditions.

**Figure S3:**
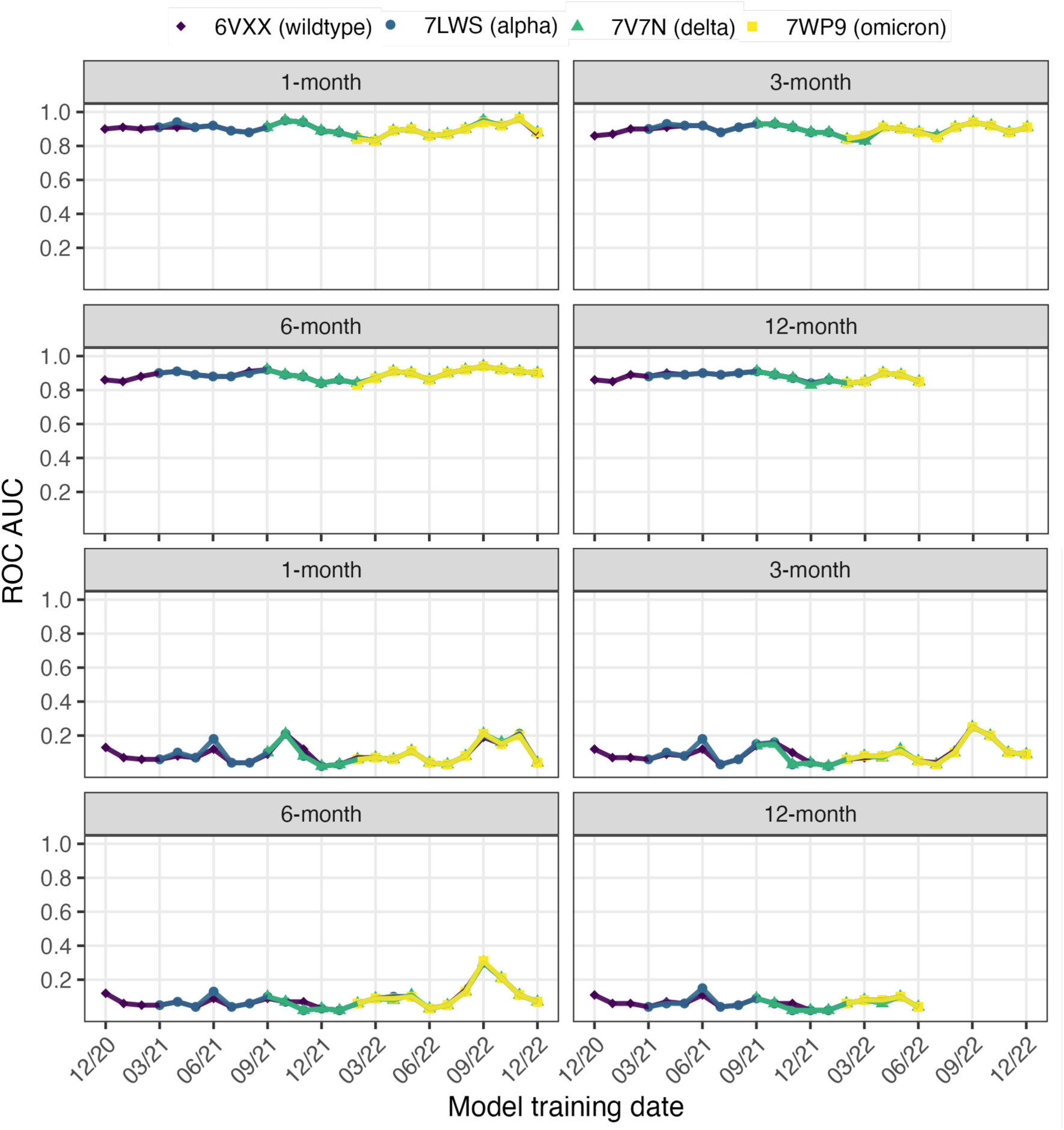
Results of training LOOS ensemble RF models at different cutoff dates with test sets sampled over different periods. AUC values are shown for all substiutions (**A**) and class-switching substiutions (**B**) curves calculated from ensemble models trained at different cutoff dates. Training data is generated by sampling of all sequences deposited 3-months before the cutoff date. Results are shown for model performance on four test sets generated from sequences deposited 1-, 3-, 6- and 12-months after the cutoff date. Four different input structures were used to calculate constraint predictors, these were 6VXX (WT), 7LWS (Alpha variant), 7V7N (Delta variant) and 7WP9 (Omicron variant). **Alt text:** Graphs showing the performance of supervised machine learning models under different training conditions, with subfigures labelled A and B.

**Figure S4:**
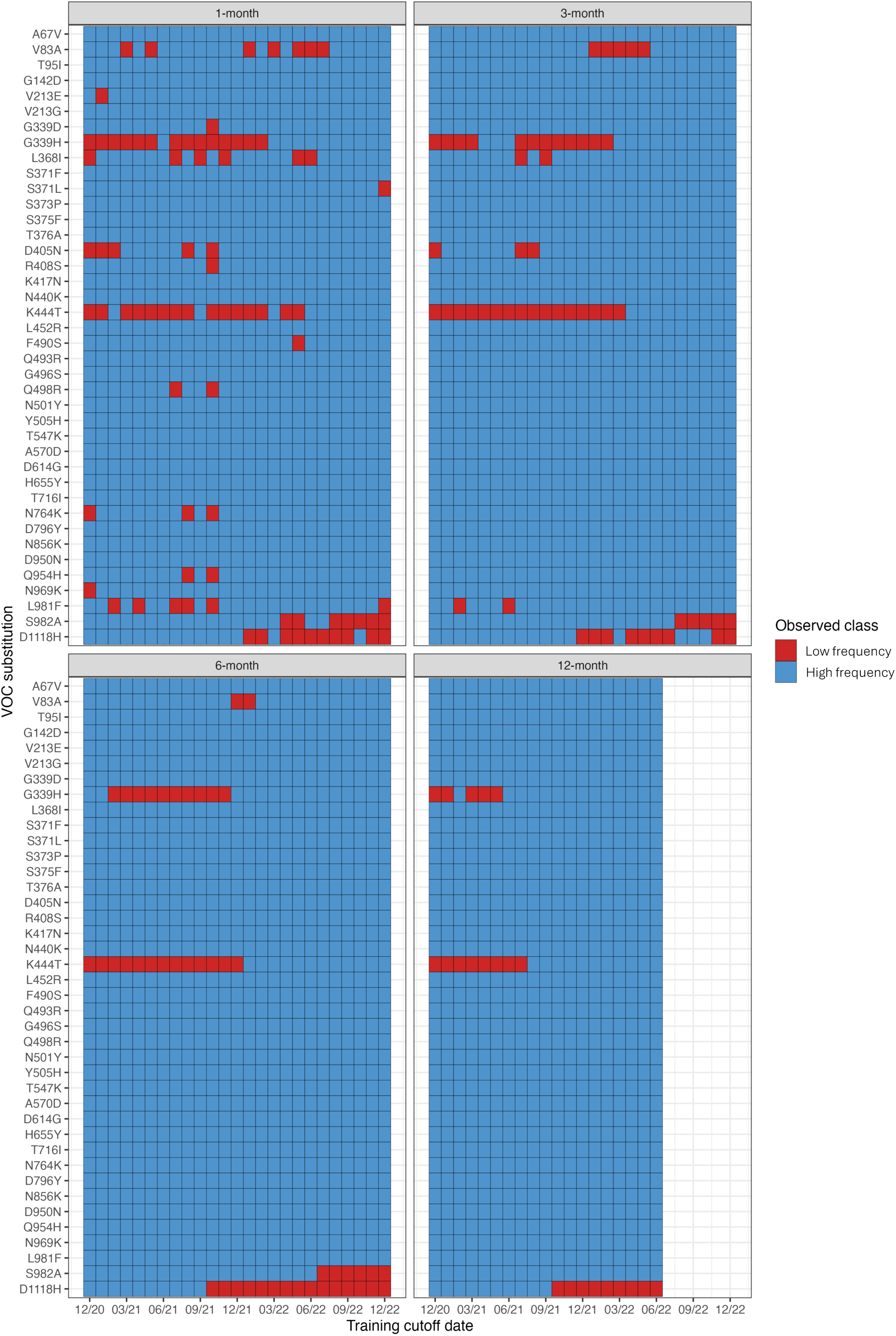
Classification of VOC substitutions over time. The observed class assigned to signature substitutions of VOCs are shown using the 1-month, 3-month, 6-month and 12-month test periods. Low frequency substitutions are shown as red, high frequency substitutions are shown as blue. **Alt text:** Heatmaps showing the classification of VOC substitutions in the SARS-CoV-2 S protein under different sampling conditions.

